# ATP citrate lyase is an essential player of the metabolic rewiring induced by PTEN loss during T-ALL development

**DOI:** 10.1101/2023.03.27.534353

**Authors:** Guillaume P. Andrieu, Guillaume Hypolite, Estelle Balducci, Caroline Costa, Els Verhoeyen, Ivan Nemazanyy, Ganna Panasyuk, Kathryn Wellen, Elizabeth Macintyre, Vahid Asnafi, Melania Tesio

## Abstract

Alterations inactivating the tumor suppressor gene *PTEN* drive the development of solid and hematological cancers, such as T-cell acute lymphoblastic leukemia (T-ALL), whereby PTEN loss defines poor-prognosis patients. We investigated the metabolic rewiring induced by PTEN loss in T-ALL, aiming at identifying novel metabolic vulnerabilities. We showed that the enzyme ATP citrate lyase (ACLY) is strictly required for the transformation of thymic immature progenitors and for the growth of human T-ALL, which remain dependent on ACLY activity even upon transformation. Whereas PTEN mutant mice all died within 17 weeks, the concomitant ACLY deletion prevented disease initiation in 70% of the animals, where it prevented the apoptosis of pre-malignant DP thymocytes by up-regulating BCL-2. Transcriptomic and metabolic analysis of primary T-ALL cells next translated our findings to the human pathology, showing that PTEN-altered T-ALL cells selectively activate ACLY and are specifically sensitive to its genetic targeting. ACLY activation thus represents a metabolic vulnerability with therapeutic potential for high-risk T-ALL patients.

## Introduction

The PI3K/AKT signaling pathway regulates essential cellular functions such as proliferation, survival, migration and metabolism. Directly opposing PI3K/AKT activation, the lipid phosphatase PTEN, (phosphatase and tensin homolog deleted on chromosome 10), fine-tunes these functions ensuring the maintenance of tissue homeostasis. *PI3K-AKT* activating mutations and/or alterations inactivating *PTEN* lead to the development of distinct human pathologies including cancer, where they are amongst the most frequent oncogenic events. This holds true not only for solid tumors but also for numerous hematological malignancies such as those arising from the transformation of thymic progenitor cells. During physiological conditions, PTEN regulates major developmental checkpoints in these cells, namely the beta selection^1^. and the positive and the negative selections^2^. During pathological conditions, instead, these cells undergo multiple genomic and non-genomic mechanisms which inactivate PTEN^3, 4^, thus driving transformation. As such, PTEN function is lost in both adult and pediatric T-cell acute lymphoblastic leukemia (T-ALL)^3, 5^. Similarly, *PTEN* alterations have been found also in pediatric T-cell lymphoblastic lymphoma (T-LBL)^6^, a closely related immature T-cell neoplasm^7^. In both these malignancies, genomic PTEN alterations have a negative prognostic impact and they identify a subgroup of patients at high-risk relapse^5, 6, 8–10^. Genomic PTEN inactivation moreover correlates with therapy resistance in T-ALL cohorts^5, 10, 11^. An unmet clinical challenge is therefore to provide novel therapies for these PTEN-altered patients with dismal outcomes.

PTEN supra-physiological expression induces a switch in the cellular metabolism, which protects the cells from the oncogenic transformation^12^. Conversely, in distinct cell types PTEN loss increases lipid biogenesis, glycolysis, protein synthesis and affects mitochondrial metabolism^13^. As in solid tumors, leukemic cells need to rewire their metabolism to face cell growth and survival. In this context, PTEN loss is likely to play an important role. We addressed this aspect in T-ALL cells, to identify novel metabolic vulnerabilities and related therapeutic targets. Combining metabolic and transcriptomic profiling, a murine T-ALL/lymphoma model and T-ALL patient-derived xenografts, we identified ATP citrate lyase (ACLY) as a crucial player in the metabolic rewiring induced upon PTEN loss during T-ALL development. This central metabolic enzyme uses coenzyme A (CoA) and mitochondrial citrate to synthesize acetyl-coenzyme A (acetyl-CoA), which is indispensable for multiple anabolic reactions such as the synthesis of fatty acids, cholesterol, and specific amino acids^14^. Activated by AKT1 in response to oncogenic signaling and cytokines stimulation^15^, ACLY moreover promotes histone acetylation in both cancer and immune cells as acetyl-CoA is also the essential cofactor of histone acetyl-transferases^16–19^. In this study, we demonstrated that ACLY is a metabolic player required for the oncogenic transformation of thymic T-cell progenitors as well as for the growth of established human T-ALL cells, where it opens a therapeutic vulnerability.

## Results

### PTEN deletion activates ATP citrate lyase in pre-leukemic thymic cells

Since PTEN regulates the cellular metabolism in distinct tissues, we hypothesized that PTEN loss could transform thymocytes by altering their cellular metabolism. To address this point, we employed a CD4Cre;PTEN^flox2^;ROSAYFP^flox2^model whereby PTEN deletion occurs in late double negative thymic progenitors and it can be monitored based on the expression of the YFP reporter. Using this model, which recapitulates PTEN-loss-induced T-ALL/lymphoma development^1, 20, 21^, we performed targeted metabolomic profiling of pre-malignant PTEN mutant thymocytes (hereby referred as PTEN^⊗^*^/^*^⊗^) (^1, 2, 21^ and data not shown), and control counterparts. As shown in **fig 1A-B**, fatty acids (palmitic, myristic, oleic, linoleic, eicosapentaenoic and docosahexaenoic acid) were among the most up-regulated metabolites in the mutant thymocytes cluster, whereas acetyl-CoA, the two-carbon donor for fatty acid synthesis, was among the most down-regulated metabolites. This was associated with a decrease in pyruvate, citrate, coenzyme A and glutamine, which paralleled an increase in glutamate and α-ketoglutarate (α-KG) (**Fig 1B-C**). These findings suggest that in PTEN mutant cells, glutamine supplies α-KG that, through reductive carboxylation, is transformed into citrate, next converted into acetyl-CoA for *de novo* lipogenesis^22^. At the crossroads of these metabolic pathways, there is the enzyme ATP citrate lyase (ACLY), which converts citrate into acetyl-CoA. Notably, ACLY is a direct target of AKT, which triggers its phosphorylation on serine 455, a post-translational modification that is indispensable to activate the enzyme^15^. Based on these data, we investigated whether ACLY is more active in PTEN mutant thymocytes. As shown in fig. 1D-E, ACLY phosphorylation on ser455 was increased in pre-malignant PTEN mutant thymocytes as compared to their control counterpart. Taken together, these results indicate that *Pten* loss activates ACLY in pre-malignant thymocytes.

**Figure 1.**
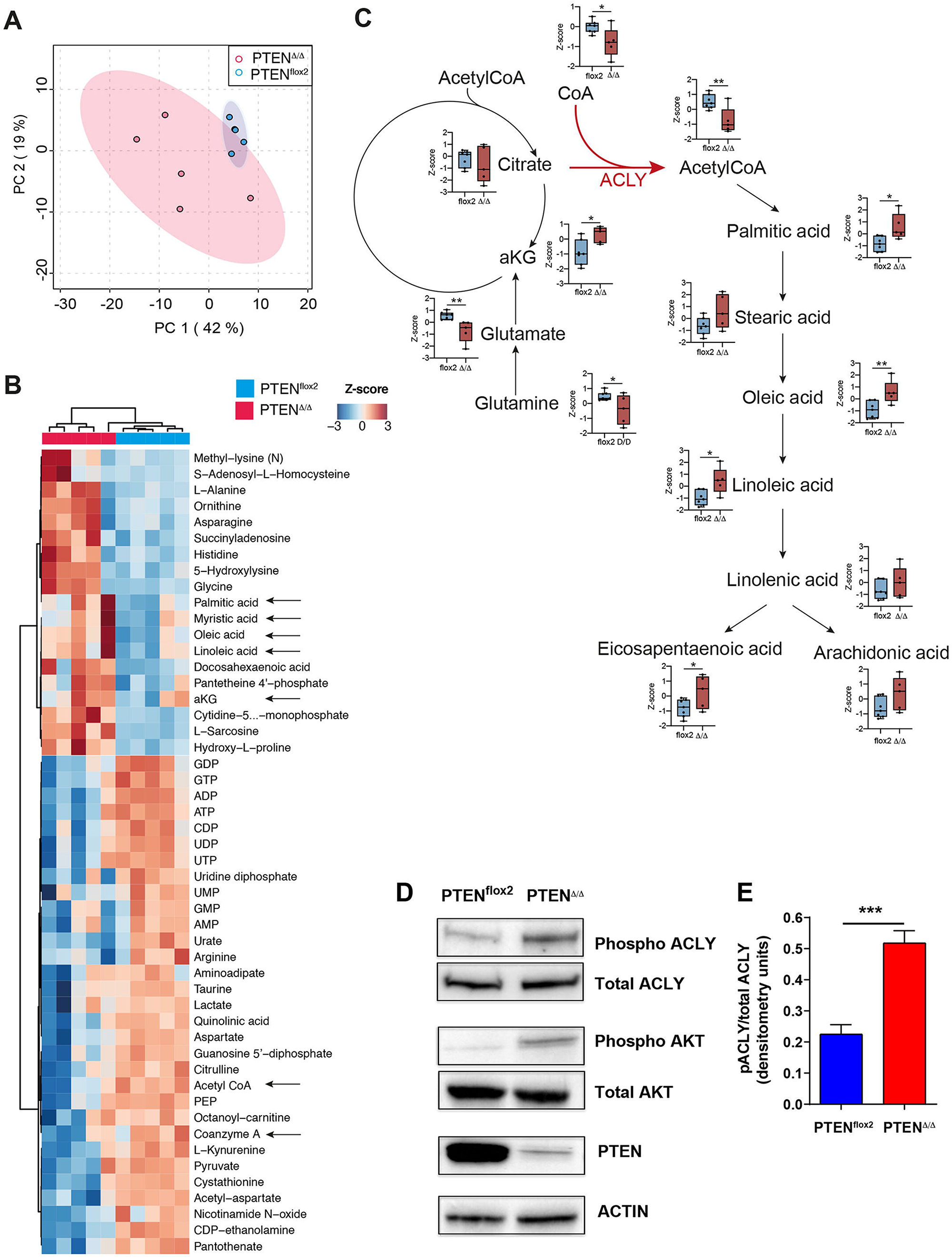
Metabolic analysis of pre-leukemic *pten* mutant thymocytes. **(A)** Principal component analysis of 4 *pten* mutant and 5 pten^flox2^ thymi issued from pre-leukemic 8 weeks old mice subjected to unsupervised metabolic analysis. **(B)** Heatmaps of the 50 most deregulated metabolites and **(C)** analysis of citrate, coenzyme A, glutamine, a-ketoglutarate, acetyl-coA and fatty acids in pre-leukemic PTEN^Δ/Δ^thymocytes or control counterparts**. (D)** Representative western blot analysis of phosphorylated ACLY (Ser455) and phoshorylated AKT (Ser473) in thymi isolated from 8 weeks old, pre-leukemic PTEN^Δ/Δ^mutant mice or PTEN^flox2^control counterparts. **(E)** Histograms depicting phosphorylated ACLY levels (ser455) in the thymi of pre-leukemic PTEN^Δ/Δ^mutant or control animals in 3 independent experiments (data are shown as average ± SE).

### Genetic ACLY ablation prevents PTEN-loss-induced T-ALL/lymphoma development

To investigate whether ACLY could promote PTEN-loss-induced leukemogenesis, we genetically ablated ACLY in PTEN mutant mice using an ACLY^flox2^model^23^ (**Suppl.** **Fig 1**). ACLY deletion abrogated the reductive carboxylation of glutamine-derived α-KG, which occurred in PTEN^null^ pre-leukemic thymocytes. Moreover it restored glutamine, citrate, coenzyme A and acetyl-CoA levels observed in wild-type mice, hence positioning ACLY as the main player in the metabolic rewiring induced by PTEN loss in pre-malignant thymocytes (**Fig 2A-C**).

**Figure 2.**
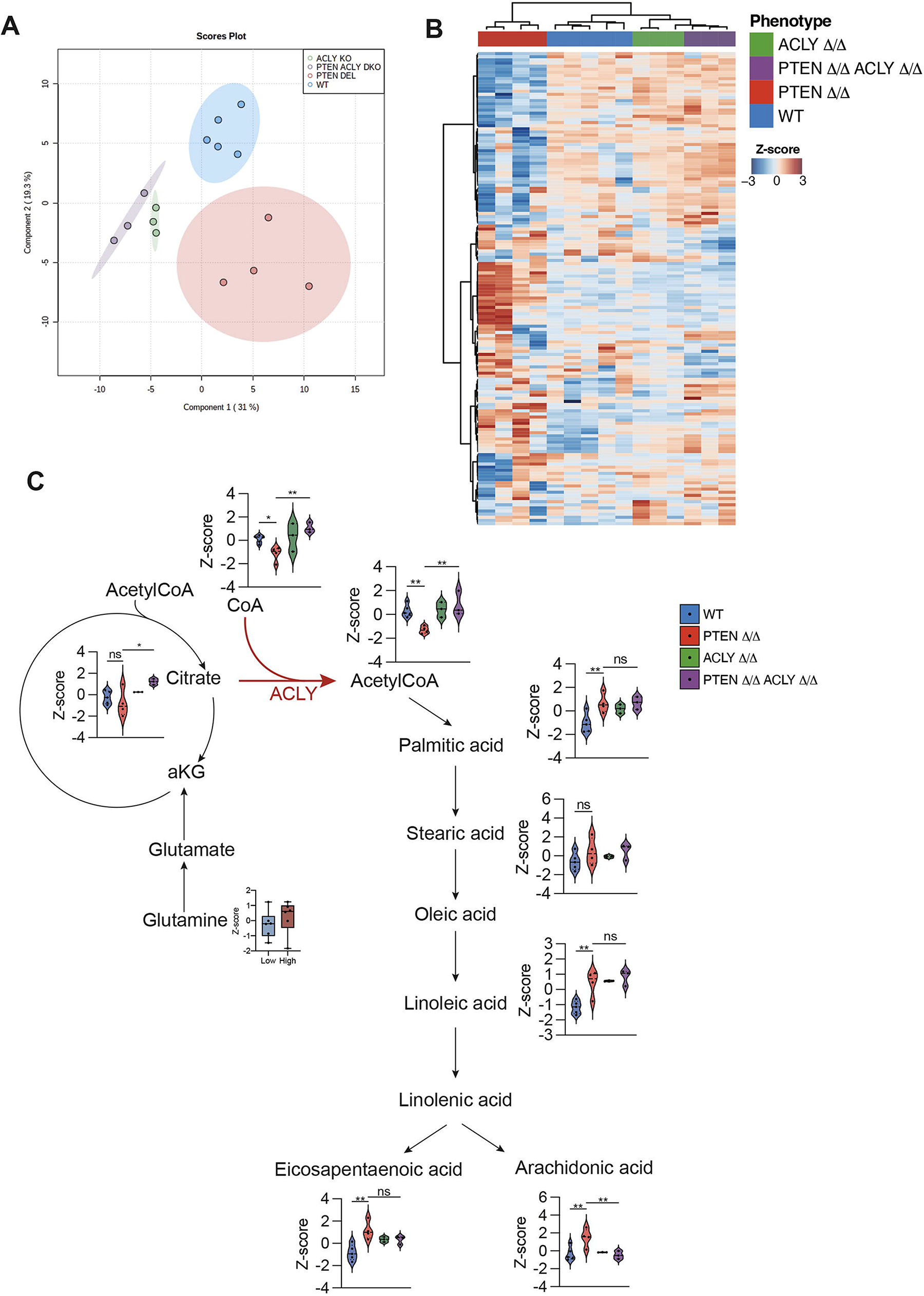
Metabolic analysis of thymi from PTEN^Δ/Δ^, ACLY^Δ/Δ^, PTEN^Δ/Δ^;ACLY^Δ/Δ^or PTEN^flox2;^ACLY^flox2^ mice. **(A)** Principal component analysis of 4 PTEN^Δ/Δ^mutant, 3 ACLY^Δ/Δ^mutant, 3 PTEN^Δ/ Δ^;ACLY^Δ/Δ^and 5 PTEN^flox2;^ACLY^flox2^ thymi issued from pre-leukemic 8 weeks old mice subjected to unsupervised metabolic analysis. **(B)** Heatmaps of the 50 most deregulated metabolites and **(C)** analysis of citrate, coenzyme A, glutamine, a-ketoglutarate, acetyl-coA and fatty acids in pre-leukemic PTEN^Δ/Δ^thymocytes or control counterparts.

Within 140 days, all PTEN^Δ/Δ^ mutant mice died of T-ALL/lymphoma (**Fig. 3A**). As previously described, these mice showed an accumulation of thymic cells bearing oligoclonal/monoclonal rearrangements of the TCRβ receptor locus and a DP TCRαβ+ phenotype (**Fig. 3B-C**). Consistent with previous findings, leukemic blasts predominantly showing a CD4 SP phenotype accumulated in blood, lymph nodes, spleen and bone marrow of PTEN mutant mice (**Fig. 3D-F** and data not shown).

**Figure 3.**
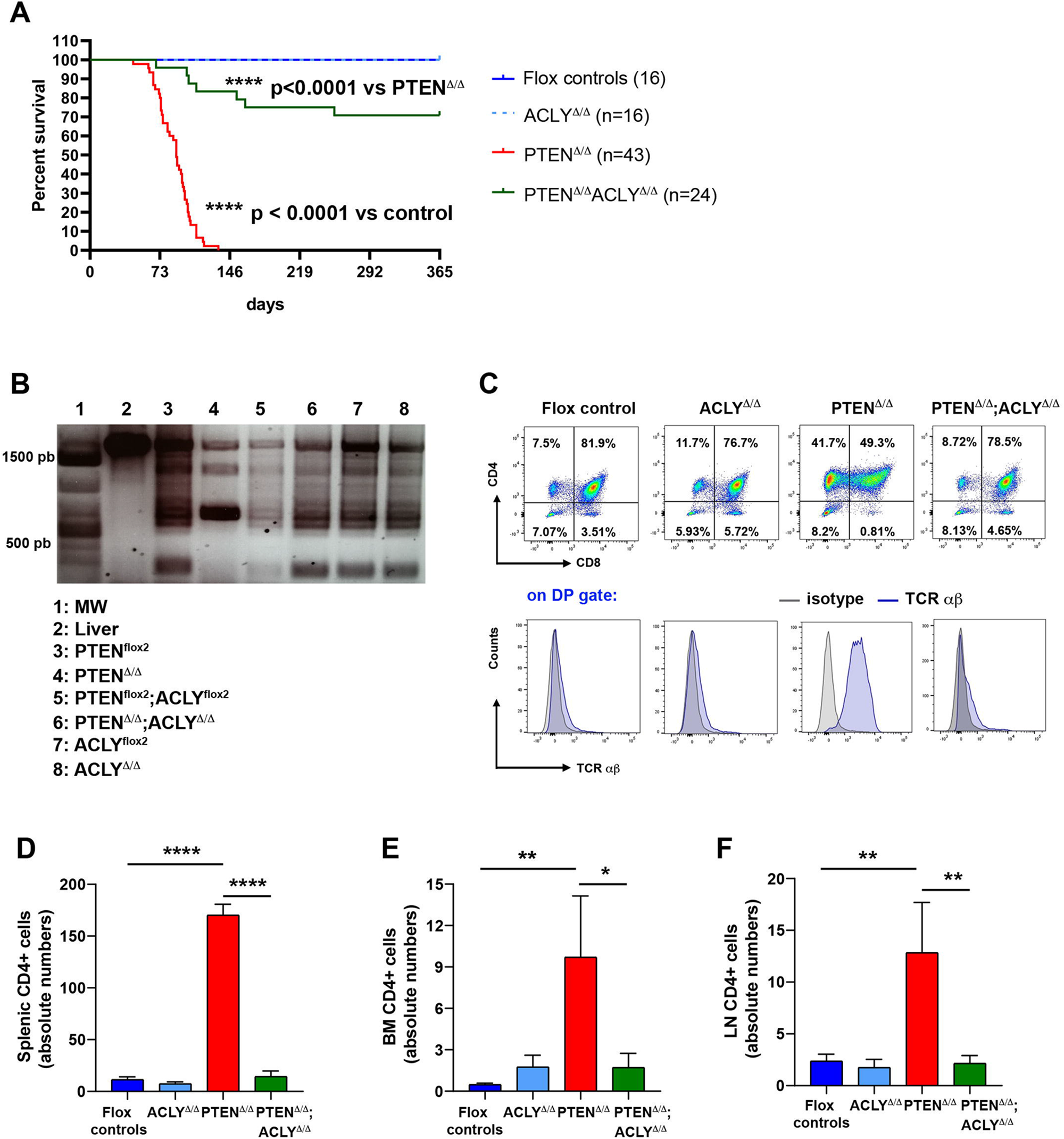
Genetic ACLY ablation prevents PTEN-loss induced T-ALL/lymphoma development. **(A)** Kaplan–Meier survival curve for PTEN^Δ/Δ^mutant mice (n=43), ACLY ^Δ/Δ^mutant (n=16), PTEN^Δ/Δ^;ACLY ^Δ/Δ^double mutant mice (n=22) and floxed control counterparts (n=40). TCRβ rearrangements **(B)** and phenotypic analysis of the thymus **(C)** in PTEN^Δ/Δ^mutant, ACLY^Δ/Δ^mutant, PTEN^Δ/Δ^;ACLY^Δ/Δ^double mutant mice or flox control counterparts. Absolute numbers of CD4+ cells in the spleen **(D)**, bone marrow **(E)** and lymph nodes **(F)** in PTEN^Δ/Δ^mutant, ACLY^Δ/Δ^mutant, PTEN^Δ/Δ^;ACLY^Δ/Δ^double mutant mice or flox control counterparts. Whereas otherwise indicated data are shown as average ± SE in at least 3 independent experiments, n=6 per group. The flox control counterparts group includes PTEN^flox2^, ACLY^flox2^and PTEN^flox2^;ACLY^flox2^mice.

In sharp contrast, only 30% of mice deleting both PTEN and ACLY (hereafter indicated as PTEN^Δ/Δ^;ACLY^Δ/Δ^) succumbed from an oligoclonal and infiltrating disease (**Fig. 3A and data not showed**), which developed with a significantly slower latency when compared to PTEN mutant mice (**Fig. 3A and suppl.** **fig 2**).

Strikingly, moreover, 70% of double mutant mice survived over one year without showing any signs of disease development. In these mice, leukemic cells were not observed in the peripheral organs or in the thymus (**Fig. 3D-F**), whereby normal polyclonal rearrangements of the TCRβ receptor locus were detected (**Fig. 3B-C**). When compared to their age-matched control counterparts, moreover, 1-year old PTEN^Δ/Δ^;ACLY^Δ/Δ^ mice presented a similar cellularity of lymphoid organs (**Suppl.** **fig 3F-I**) as well as a similar distribution of DN, DP and SP thymic progenitors (**Suppl.** **fig 3E**). ACLY deletion *per se* did not alter mice survival (**Fig. 3A**) nor the cellularity of lymphoid organs (**Suppl.** **fig 4A-D**).

**Figure 4.**
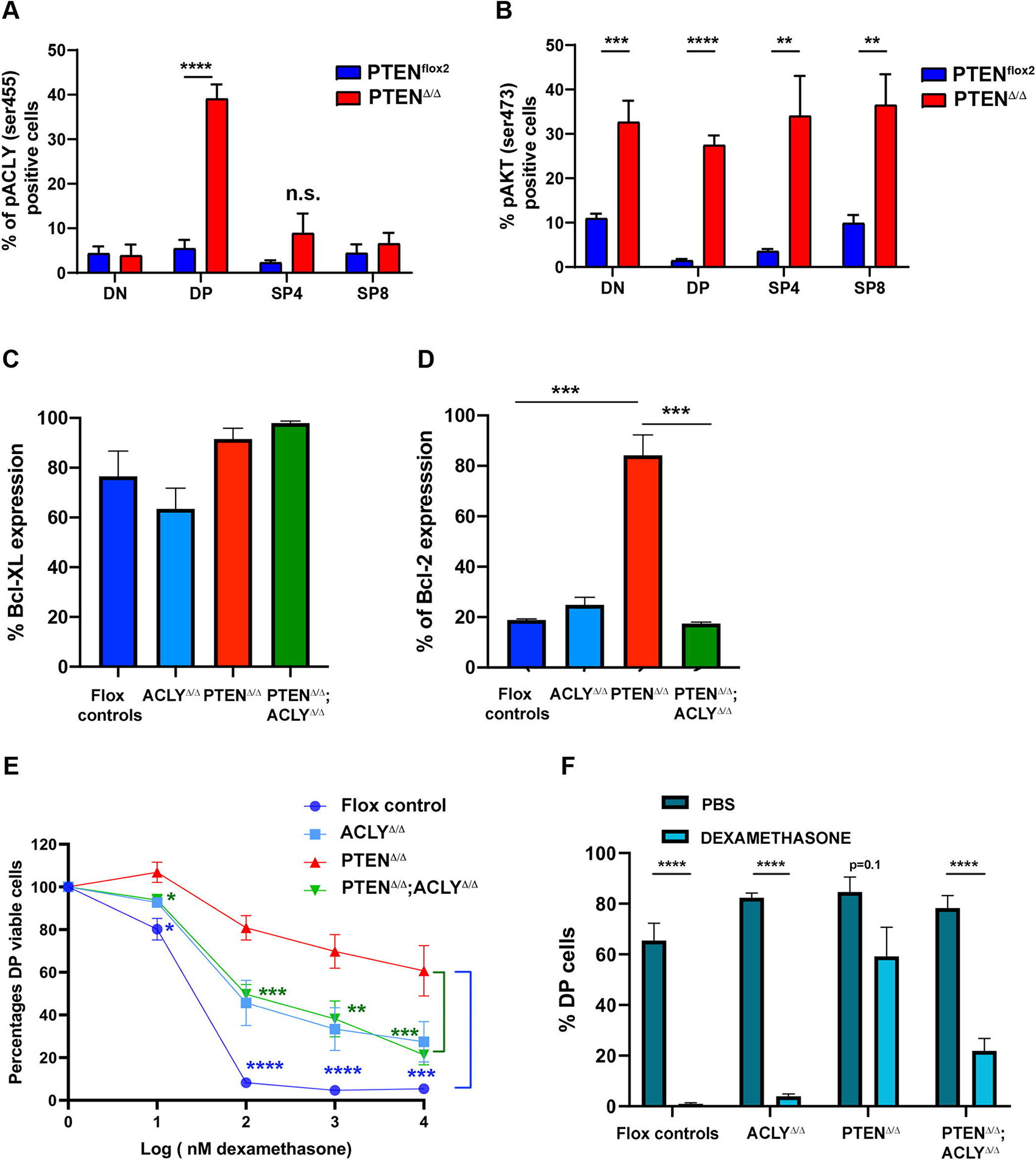
ACLY ablation abrogates apoptosis resistance in DP PTEN null thymic progenitors. **(A)** Phospho AKT (ser473) and **(B)** and phospho ACLY (ser455) levels on DN, DP, SP4 and SP8 from pre-leukemic PTEN^Δ/Δ^mutant or PTEN^flox2^control counterparts. In mutant mice, which carried the YFP^flox2^reporter, DN, DP, SP4 and SP8 cells are gated on YFP+ cells. Levels of Bcl-xL **(C)** and BCL-2 **(D)** in DP cells from PTEN^Δ/Δ^mutant, ACLY^Δ/Δ^mutant, PTEN^Δ/Δ^;ACLY^Δ/Δ^double mutant mice or flox control animals **(E)** Sensitivity of DP cells from PTEN^Δ/Δ^mutant, ACLY^Δ/Δ^mutant, PTEN^Δ/Δ^;ACLY^Δ/Δ^double mutant mice or flox control animals to dexamethasone treatment ex vivo. **(F)** Percentages of DP cells in PTEN^Δ/Δ^mutant, ACLY^Δ/Δ^mutant, PTEN^Δ/Δ^;ACLY^Δ/Δ^double mutant mice or flox control animals receiving PBS or dexamethasone.

Taken collectively, these results demonstrate that ACLY plays a crucial role in PTEN-loss-induced T-ALL development by promoting disease initiation.

### Genetic ACLY ablation prevents PTEN-loss-induced resistance to apoptosis

As previously shown, in PTEN mutant mice the disease originates from apoptosis-resistant pre-leukemic DP cells which are transformed into leukemia-initiating cells following the acquisition of additional oncogenic events^2, 24^. Being ACLY required to initiate the disease in PTEN mutant mice, we next addressed whether, on a mechanistic level, this could results from ACLY ability to promote apoptosis resistance in PTEN null pre-leukemic cells.

To investigate this hypothesis, we first analyzed whether ACLY activation selectively occurs in pre-leukemic PTEN null DP cells. As shown in **Fig. 4A-B**, despite PTEN ablation activated AKT in double negative (DN), double positive (DP), CD4 single positive (SP4) and CD8 single positive (SP8) thymic progenitors, a significant increase in phospho ser455 ACLY levels was only observed in mutant DP cells, thus demonstrating that PTEN loss activates ACLY exclusively in the cells of origin of the disease. We next analyzed whether ACLY activation in these cells modulates the expression of anti-apoptotic proteins. As shown in **Fig 4C**, we did not observe major differences in the expression of BCL-XL, the main anti-apoptotic protein in DP thymic cells^25, 26^. Yet, in these cells, which developmentally express very low BCL-2 during physiological conditions^27, 28^, PTEN loss induced a dramatic increase in BCL-2 expression. Interestingly, this was prevented by ACLY deletion (**Fig. 4D**). These data, showing that ACLY is instrumental to up-regulate BCL-2 in pre-leukemic PTEN null DP cells, next prompted us to functionally examine whether its deletion prevents PTEN-loss induced resistance to apoptosis in these cells.

As shown in **Fig. 4E**, whereas DP cells sorted from PTEN mutant mice did not undergo apoptosis when treated *ex-vivo* with dexamethasone, DP cells sorted from PTEN^Δ/Δ^;ACLY^Δ/Δ^ mice were highly sensitive to dexamethasone-induced apoptosis. In line with this, ACLY deletion in PTEN null mice reduced the *in vivo* dexamethasone-resistance of pre-leukemic DP cells. (**Fig. 4F**). ACLY deletion *per se* increased the dexamethasone resistance of DP cells as compared to control conditions both ex-vivo and in vivo (**Fig. 4E-F**). Taken together, these data suggest that ACLY activation upon PTEN loss promotes the resistance of pre-leukemic DP progenitors to apoptotic stimuli, thus potentially explaining its role in promoting disease initiation.

### Human PTEN null T-ALL cells modulate ACLY activity

We next investigated whether our findings could be recapitulated in the human setting. To do so, we first investigated whether PTEN silencing in a panel of T-ALL PTEN germ-line cell lines was able to activate ACLY. As shown in **Fig. 5A-B**, the transduction of five distinct cell lines with a lentiviral vector carrying a shRNA PTEN significantly increased ACLY phosphorylation on ser455 and acetyl-CoA levels, consistent with an AKT1-mediated activation of the enzyme in these cells (**Fig. 5A**). In line with these findings, ACLY targeting by a shRNA or a selective pharmacological inhibitor prevented the growth of RPMI and DND41 cells (**Fig 5C**). Noteworthy, these effects selectively occurred in PTEN null cells but not in PTEN germ-line cells, thus suggesting that only PTEN-altered cells are specifically vulnerable to ACLY inhibition. To further confirm these findings and translate them to the patients setting, we performed a metabolic analysis of 8 patients derived xenografts (PDX). This included 3 patients with PTEN abnormalities, 1 patient with PI3KCA mutations and 4 patients with a germ line status for PTEN, PI3KCA, PI3KR1 and AKT1 (**Table 1**). Consistent with the reported genomic alterations, with the exception of one PTEN altered case, patients bearing PTEN/PI3KCA alterations displayed significantly higher levels of phosphorylated AKT (ser347) (**Table 1**). This mirrored the clustering identified by the principal component analysis (**Fig. 6A**). Paralleling what observed in the murine model, patients bearing genomic abnormalities in the PI3K/PTEN pathway displayed significantly lower levels of coenzyme A and acetyl-coA and they showed a de-regulated Krebs cycle (**Fig 6B-D**). In line with our murine models, PTEN altered cases presented a metabolic profile polarized toward the synthesis of fatty acids such as palmitic and stearic acids (**Fig 6C-E**). Enrichment analyses revealed the prevalence of tRNA synthesis, amino acids and unsaturated fatty acids, consistent with acetyl-coA consumption (**Fig. 6D** and **Suppl. Fig. 5**). Hence, our results suggest that PTEN loss of function shifts the blasts metabolism towards fatty acid synthesis notably by modulating ACLY. To further investigate this aspect, we carried out a transcriptomic analysis of 155 primary T-ALL by RNA sequencing. Differential expression analysis of 4,871 metabolic genes revealed that most of the PTEN-altered cases compose a cluster (**Fig. 7A**). In line with our metabolomic analysis, the over-representation analysis of deregulated metabolic genes in PTEN-altered cases revealed an up-regulation of fatty acids and long-chain fatty acyl-CoA esters synthesis, and a down-regulation of CoA synthesis (**Fig. 7B**). We next stratified primary samples based on inferred PI3K activity as determined by PROGENy. This analysis confirmed that an elevated PI3K signaling is associated with the up-regulation of fatty acid synthesis and the TCA cycle (**Fig. 7C-E**). A strong PI3K activity also correlated with the enrichment in a cholesterol signature (**Fig. 7F**) and an increased cholesterol synthesis was observed in PTEN-altered PDX-derived blasts (**Fig. 7G**). Taken together, these analyses demonstrated that ACLY activation is at the center of a crucial metabolic rewiring induced by PTEN loss in T-ALL cells, which is likely to provide fatty acids and cholesterol necessary to their growth.

**Figure 5.**
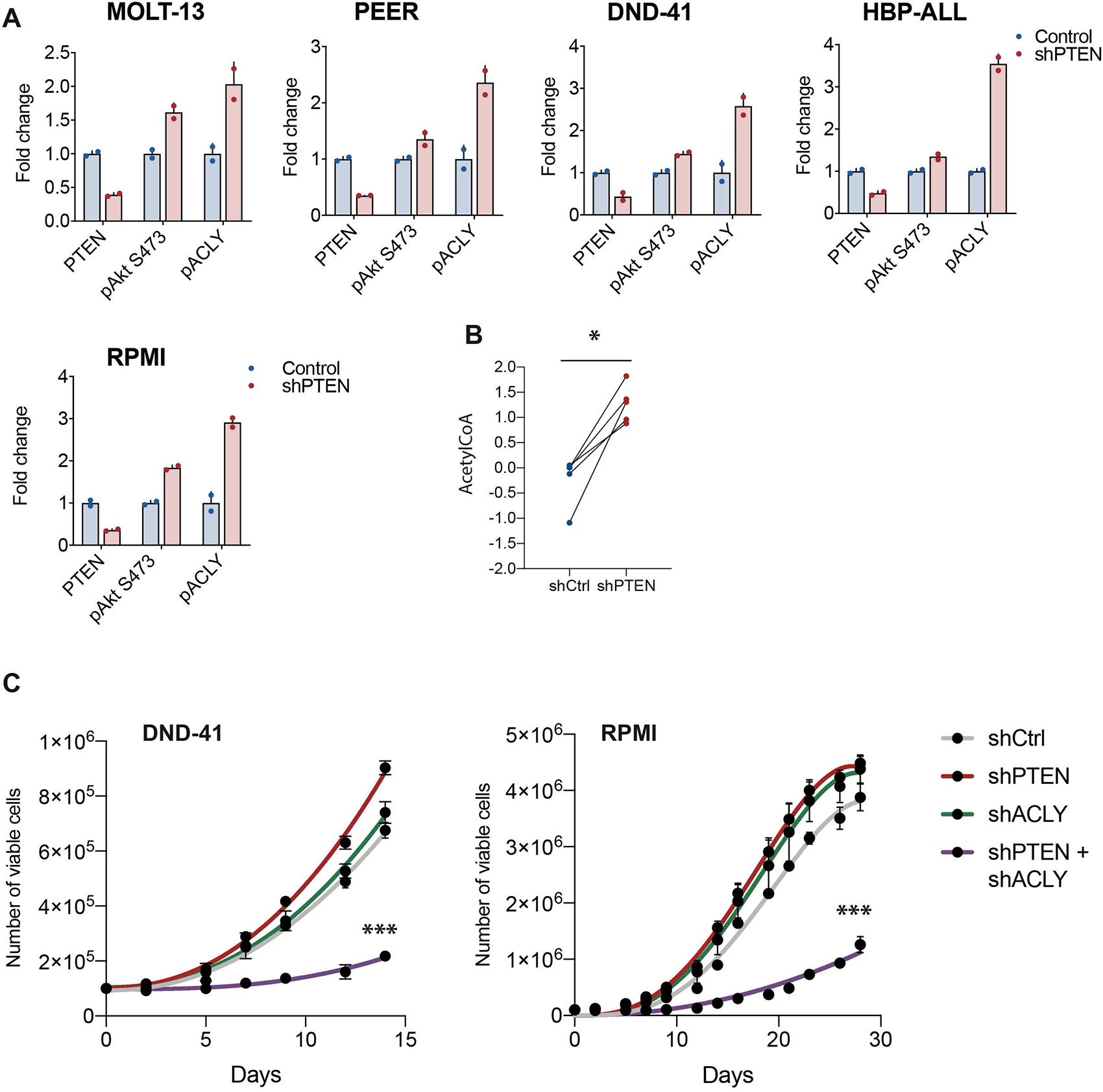
PTEN loss activates ACLY in human T-ALL cells to sustain cell growth. PTEN, pAKTser473 and pACLYser455 levels **(A)** and acetyl-coA levels **(B)** in a panel of human T-ALL cell lines transduced with a lentiviral vector carrying an shRNA PTEN or a control shRNA. **(C)** Growth of DND41 and RPMI cells transduced with an shRNA PTEN, an shRNA ACLY or the combination of the two and relative control shRNA.

**Figure 6.**
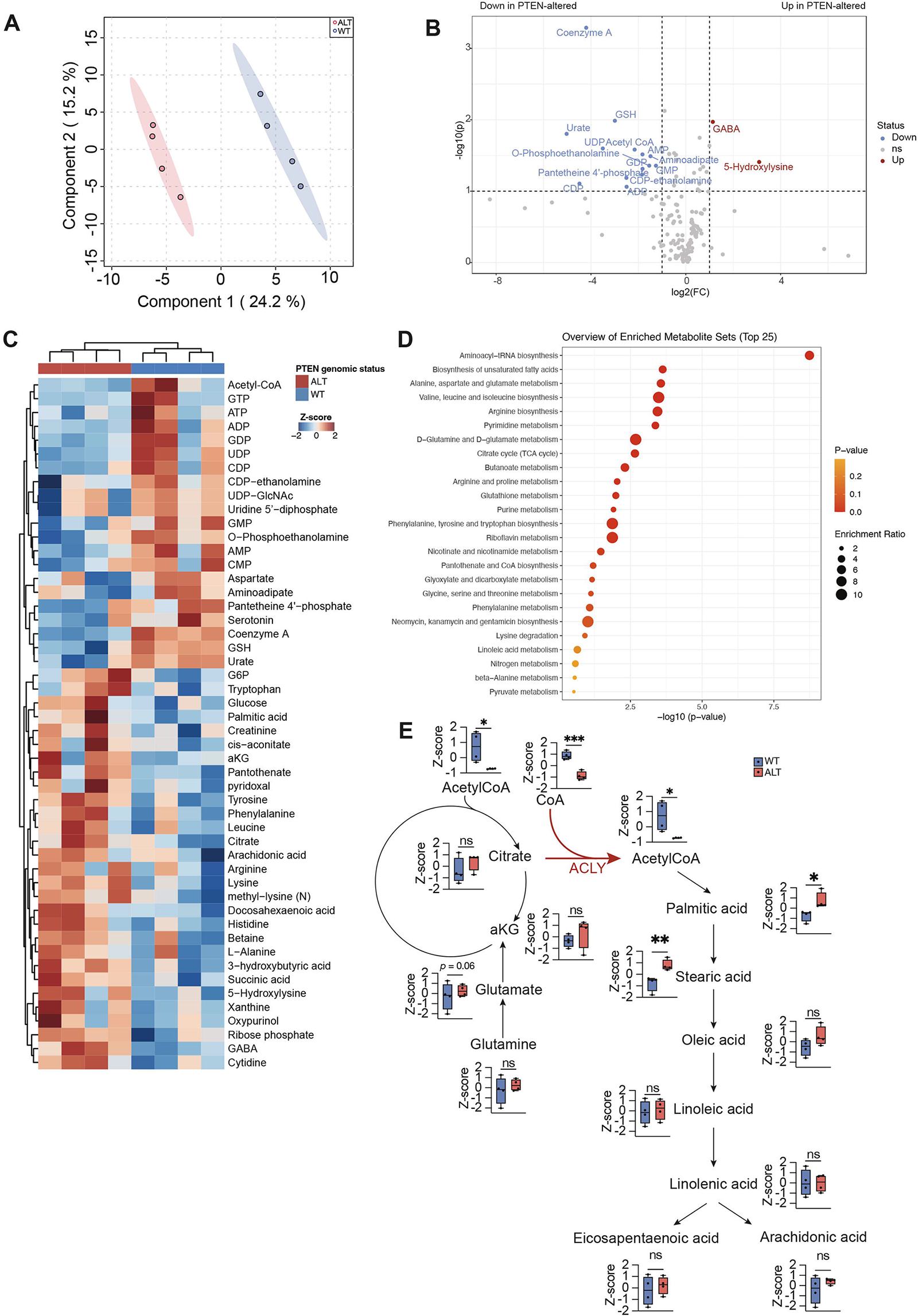
Targeted metabolic analysis of patents derived T-ALL cells. **(A)** Principal component analysis of 8 PDX which underwent targeted metabolomic analysis. **(B)** Most up-regulated and down-regulated metabolites in PTEN-altered cases. **C)** Heatmaps of the most 50 deregulated metabolites and **(D)** pathways enriched in PTEN-altered patients. **(E)** Metabolic analysis of citrate, coenzyme A, glutamine, a-ketoglutarate, acetyl-CoA and fatty acids in PTEN-altered cells

**Figure 7.**
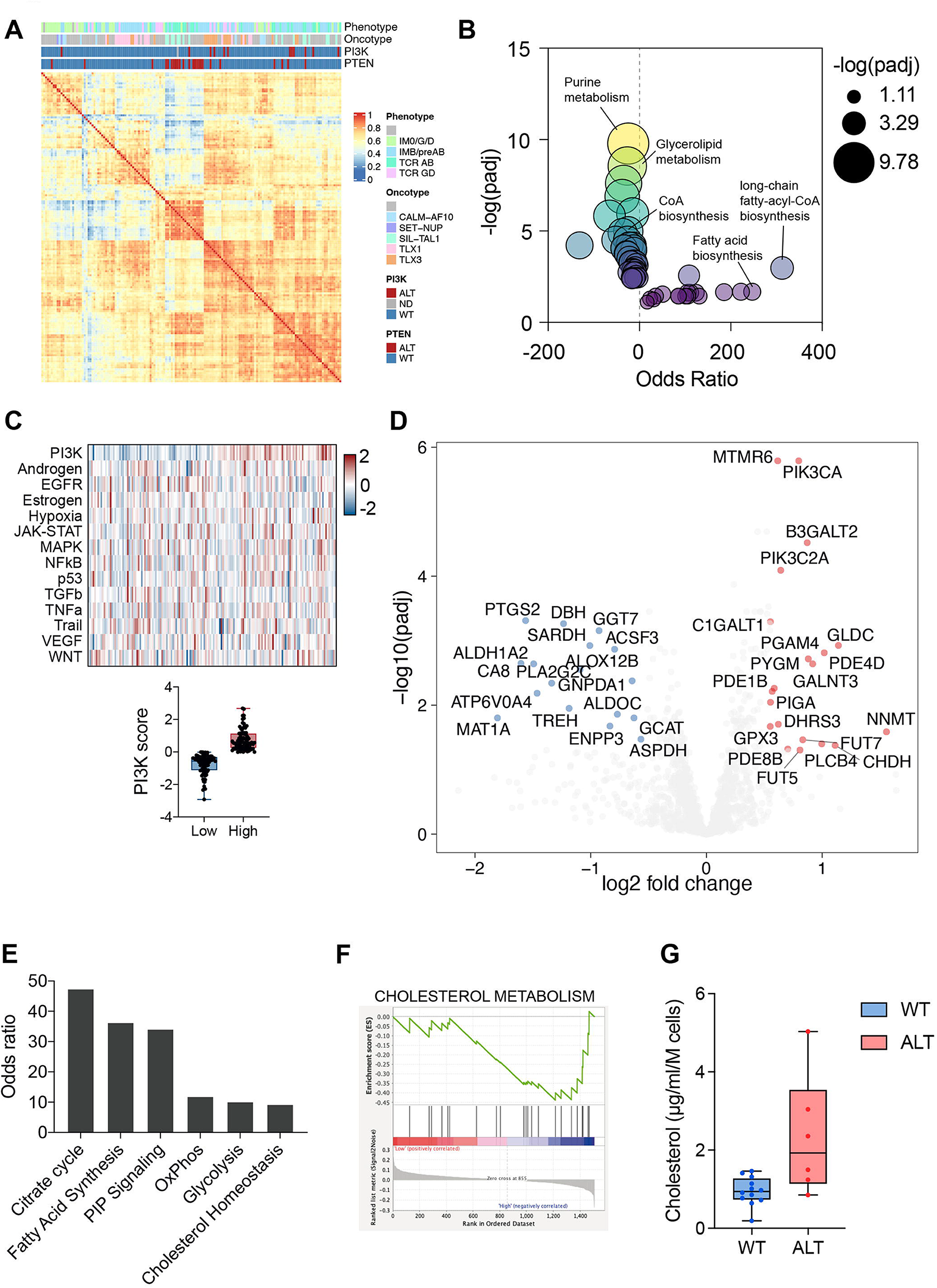
Transcriptomic analyses reveal the characteristics of PI3K signaling-altered T-ALL patients. **(A)** Metabolic transcriptome-based clustering of 155 patients reveals that PI3K signaling alterations define two homogeneous groups of patients sharing similar oncogenetic features. **(B)** Pathway enrichment analyses based on the differential metabolic gene expression of PTEN-altered (n=33) vs wild-type (n=122) patients determined using EnrichR. **(C)** PROGENy score computation from whole transcriptomics data from 155 patients. The median score of the series was used to stratify patients into high and low-score groups. **(D)** Volcano plot depicting the differentially expressed metabolic genes in high vs low PI3K score. **(E-F)** Enriched metabolic gene sets enrichment in high vs low PI3K score as determined by GSEA. **(G)** Intracellular levels of cholesterol measured in WT or altered PI3K signaling T-ALL PDX blasts.

**Table 1.** PTEN, PI3K3A, PI3R1, AKT1 status and phospho AKT levels in 8 PDX which underwent metabolomic analysis.

### PTEN-altered T-ALL cells are specifically sensitive to ACLY targeting

To further corroborate our findings, we next analyzed phospho ACLY (ser455) levels in additional 14 PDX-derived blasts T-ALL cells bearing *PTEN/PI3KCA* alterations and showing active AKT signaling. As shown in **Fig. 8A**, these patients displayed higher ACLY phosphorylation, consistent with an AKT1-mediated activation of ACLY. Based on these results, we next silenced ACLY in five different PDX-derived blasts by transducing these cells with a baboon-pseudotyped lentiviral vector carrying an ACLY shRNA ACLY or a control shRNA^29^. Remarkably, in four out of five PTEN-altered PDX which we analyzed, ACLY silencing had a pronounced effect on their *in vitro* viability already 48 hours after the transduction. As shown in **Fig. 8B**, at this time point already a few viable GFP-positive cells were left among those transduced with the shRNA ACLY-carrying vector. In the fifth PDX derived sample, ACLY targeting did not affect the cellular viability ex-vivo, prompting us to examine its effect in vivo. We performed therefore a competitive transplantation assay, whereby transduced cells were allowed to compete with non-transduced cells in a 20:80 ratio following their injection into NSG mice. As shown in **Fig. 8C**, PDX-derived blasts lacking ACLY were out-competed by non-transduced cells, whereas GFP control cells were not. We next investigated whether ACLY represents a metabolic vulnerability specific to PTEN-altered T-ALL cases. To do so, we genetically silenced either PTEN or ACLY or both of them in PDX-derived cells displaying a germ line PTEN status. As shown in **Fig. 8D**, ACLY silencing reduced the growth of these primary cells only when PTEN was concomitantly silenced. Taken together, these results indicate that ACLY is indispensable to promoting the survival of PTEN altered T-ALL cells, where it represents a specific metabolic vulnerability.

**Figure 8.**
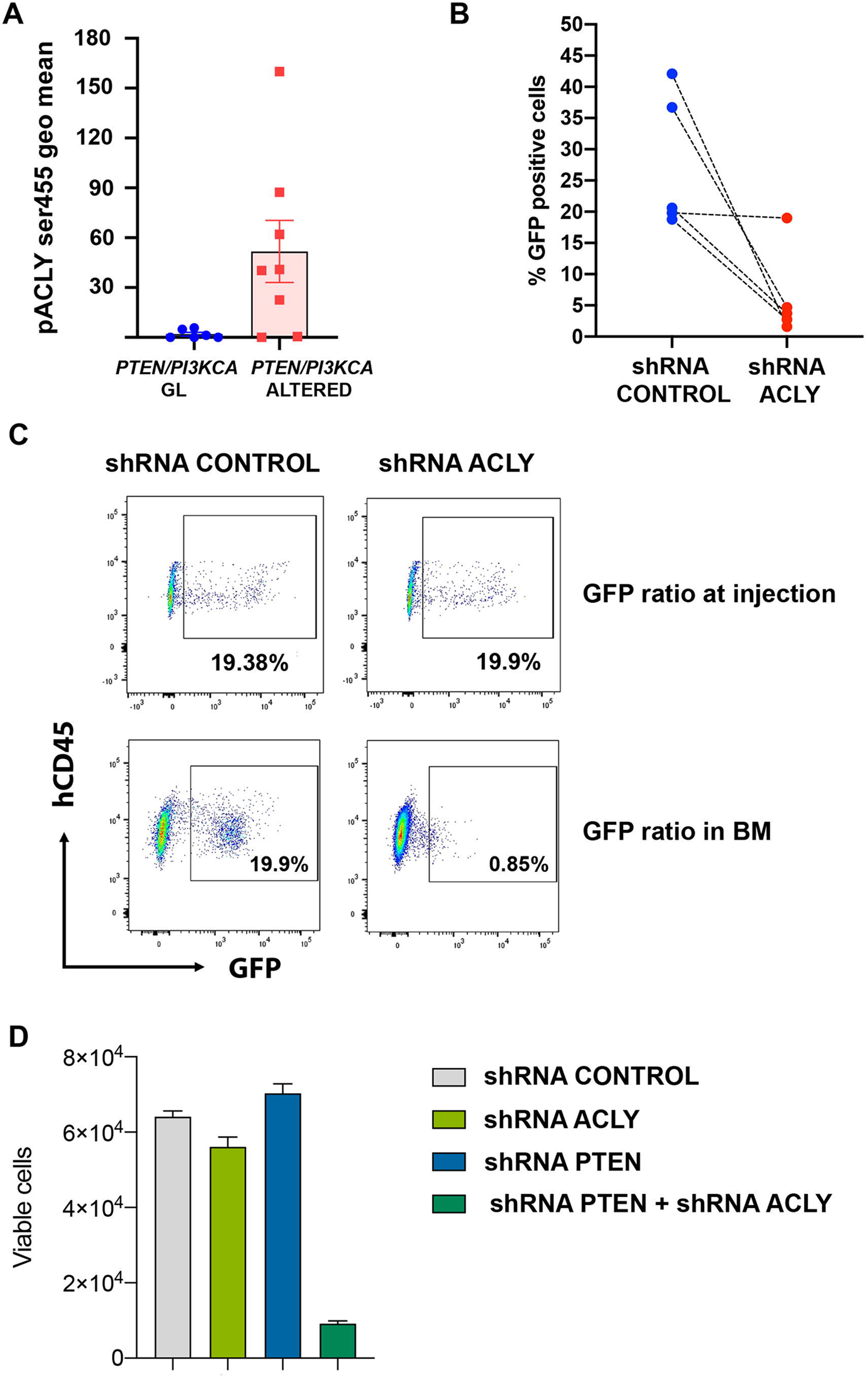
ACLY silencing in primary PTEN null T-ALL cells selectively affects their viability. **(A)** Phospho ACLY ser455 levels in 8 *PTEN/PI3KCA* altered and 6 germ line PDX-derived T-ALL blasts. (**B)** Percentages of GFP+PTEN altered PDX-derived blasts which remained viable 48 hours post-transduction with lentivirus vectors carrying shRNA ACLY or a control construct. **(C)** Competitive transplantation assay, whereby PDX-derived cells were transduced with a shRNA ACLY or a shRNA control and allowed to compete with non-transduced cells in a 20:80 ratio following their injection in NSG mice. The relative ratio of transduced and not transduced cells in the bone marrow of NSG mice injected with either shRNA ACLY or shRNA control transduced cells is shown. **(D)** Number of GFP+ PTEN germ-line PDX-derived blasts which remained viable 72 hours post-transduction with lentivirus vectors carrying a shRNA ACLY, a shRNA PTEN, the combination of the two shRNA or relative control constructs.

## Discussion

We demonstrated that ACLY is a metabolic player not only required for the oncogenic transformation of thymic T-cell progenitors but also for maintaining an established human T-ALL, which represents an important metabolic vulnerability.

Combining metabolomics and a murine T-ALL/lymphoma model, we showed that PTEN loss activates ACLY in a selective manner in pre-malignant DP thymocytes and that this is instrumental to drive disease initiation. From a mechanistic point of view, we show that this arises, at least in part, from an ACLY-mediated regulation of apoptosis resistance in these cells, which are the cells-of-origin of the disease^24, 30^. Whereas during physiological T-cell development, DP progenitors down-regulate BCL-2 in favor of BCL-XL, we show that in pre-leukemic conditions, PTEN loss does not affect BCL-XL expression but rather induces supra-physiological levels of BCL-2 and that this occurs via ACLY activation. This may imply an epigenetic regulation of BCL-2 expression as as the ACLY-derived acetyl-coA serves as a HAT cofactor to acetylate histones and activate gene transcription^16^. Being the sole donor of acetyl groups for acetylation, acetyl-CoA also critically regulates protein acetylation on N-terminal residues (N-terminal acetylation) and on lysine residues (Nε acetylation), a post-translational modification which affects proteins stability as well as the cross-talk with other post-translational modifications^13^. ACLY may thus additionally regulate BCL-2 levels on a post-translational level favoring its acetylation. In line with this scenario, acetyl-CoA was shown to regulate apoptotic sensitivity by promoting BCL-xL N-alpha-acetylation^32^ and an interplay between protein acetylation and ubiquitination has been recently shown to control the stability of the anti-apoptotic protein MCL-1^33^. Whereas further studies will be necessary to fully dissect how exactly ACLY couples PTEN loss and BCL-2 up-regulation to promote apoptosis resistance, our data pointing out a role for ACLY in glucocorticoid-induced apoptosis resistance are intriguing as *PI3K/PTEN* alterations are associated with glucocorticoid resistance in clinical settings^5, 8, 11^.

As our translational investigations point-out, ACLY role in T-ALL development relates not only the oncogenic transformation of immature thymic progenitors but also the maintenance of an established leukemia, which remains dependent on ACLY activity even upon transformation.

In this context, ACLY anabolic functions are likely to play an important role since in *PTEN/PI3K* altered cells metabolic, genetic and transcriptomic profiling of human T-ALL cells evidenced an increased need for ACLY to supply acetyl-coA for fatty acids and cholesterol biosynthesis. These findings, which recapitulate the results obtained with the murine model, are in line with previous data highlighting a role for cholesterol biosynthesis in the growth of immature ETP-ALL cells^34^. Consistent with its selective activation in PTEN-altered patients, ACLY specifically sustains the growth of T-ALL cells that underwent PTEN loss. As evidenced in both cell lines and primary T-ALL, only T-ALL cells deprived of PTEN were sensitive to ACLY silencing.

Highlight a specific metabolic vulnerability linked to PTEN/PI3K genetic alterations, our data suggest that high-risk T-ALL patients bearing these abnormalities may benefit from pharmacological ACLY targeting. Albeit ACLY inhibitors have been under development for a few years, the only FDA-approved ACLY inhibitor (Bempedoic acid) also stimulates AMPK activity and thus lacks selectivity. Conversely, the ACLY inhibitors which showed good specificity did not reach clinical studies, as in the case of SB-204490 a poor tissue-specific distribution in humans has limited them^36^. In light of this, the results of out study holds the promises of boosting the development of novel specific ACLY inhibitors suitable for clinical intervention.

In conclusion, our study revealed that PTEN loss defines a specific metabolic rewiring in T-ALL cells, whereby ACLY is a major player as it favors not only the transformation of thymic progenitor cells but also the growth of established human leukemias. Primary T-ALL cells dependency on ACLY activity thus represents a novel metabolic vulnerability with therapeutic potential.

## Material and methods

### Mice

PTENflox/flox mice (B6.129S4-Ptentm1Hwu/J), generated as previously described (REF), were crossed with CD4Cre mice (B6.Cg-Tg(Cd4-cre)1Cwi/BfluJ) and the reporter line ROSAYFPflox/flox (B6.129X1-Gt(ROSA)26Sortm1(EYFP)Cos/J) to obtain CD4Cre;PTENflox2,ROSAYFPflox2 mice. ACLYflox/flox mice (Aclytm1.1Welk/ Mmjax), generated as previously described ^23^, were crossed with CD4Cre;PTENflox2,ROSAYFPflox2 mice to obtain double mutant CD4Cre;PTENflox2;ACLYflox2;ROSAYFPflox2, the single mutant mice CD4Cre;ACLYflox2;ROSAYFPflox2 and CD4Cre;PTENflox2,ROSAYFPflox2 mice and relatve control counterparts (respectively PTENflox2;ACLYflox2;ROSAYFPflox2, ACLYflox2;ROSAYFPflox2 and PTENflox2,ROSAYFPflox2 mice). All mice were purchased from Jackson laboratory and showed a full C57/Bl6J genetic background as evidenced by SNP analysis. These mice were maintained under SPF (Specific Pathogen Free) conditions. NOD Scid Gamma Mice ( NOD. Cg-Prkdc <SCID>Il2rg<TM1WJL>/SzJ) were purchased from Charles River Laboratories and use for transplantation assays and xenograft assays. These mice were maintained under SOPF (Specific and Opportunistic Pathogen Free) conditions. All experiments were performed according to procedures approved by the Ethical committee of Paris Descartes University and the Ministry of Higher Education, Research and Innovation (APAFIS#25489-2019091313386161 v3).

### Lymphoid organs isolation

To collect BM cells, mice femora, tibiae and hips were dissected and crushed using a mortar and pestle, and cell suspensions were filtered before further use. Mouse spleens, thymi and lymph nodes were isolated and crushed using the bottom of a syringe punch and filtered to obtain cell suspensions. Blood cells were collected from cheek vein bleedings into tubes containing 10,0000 U/ml heparin (Sigma-Aldrich), and peripheral red blood cells were lysed using ACK lysis buffer (Thermo Fisher Scientific). Organ cellularity was determined by an automated CASY cell counter (OMNI Life Science 5651736).

### Murine thymocytes cultures

Thymocytes harvested from the mice were cultivated in home-made thymocytes culture medium (250ml water, 2,5g a-MEM in powder, 7,35ml NaHCO3, 52.5ml serum hyclone, 3.1ml l-glutamine, 3.1ml pen/strept, 316µl 2-mercaptoethanol 50mM, 3.16ml HEPES 1M, 3.16ml sodium pyruvate 100x, 63µl gentamicin 50mg/m) with mouse cytokines mIL-7 (5ng/ml) (RnD Systems 407-ML-005) and mFLT3 (5ng/ml) (RnD Systems 427-ML-005) and treated with Dexamethasone (0, 10nM, 50nM, 100nM, 1uM, 10uM) for1 8 hours.

### Analysis of murine TCRb rearrangements

DNA was extracted from sorted DP thymocytes using the DNeasy Blood and Tissue Kit (Qiagen). As previously described (REFs), rearranged TCRβ regions were amplified using Vβ2 forward (5′-GTAGGCACCTGTGGGGAAGAAACT-3′) or Dβ2 forward (5′-GGGTCACTGATACGGAGCTG -3′) primers and Jβ2 reverse primer (5′-TGAGAGCTGTCTCCTACTATCGATT -3′) primer.

### Patients

Diagnostic peripheral blood or bone marrow samples from adult T-ALL patients were analyzed after informed consent was obtained at diagnosis according to the Declaration of Helsinki. Patients were included in the GRAALL03/05 trials, both were registered at <u>ClinicalTrials.gov</u> (GRAALL-2003, NCT00222027; GRAALL-2005, NCT 00327678). The screening of oncogenetic abnomalies as the immunophenotyping of the samples were performed as previously described (Andrieu GP et al Blood 2021).

### RNA sequencing of patient samples

Total RNA extraction was performed using the RNeasy Mini Kit (Qiagen) and converted into cDNA using SuperScript III Reverse transcription (Thermo Fisher Scientific). Libraries were prepared using the SureSelect XT HS2 RNA System (Agilent) and sequenced on a NovaSeq (Illumina). Differential expression and pathway enrichment analyses were performed as previously described^36^. For metabolic-specific analyses, the entire dataset was reduced to metabolic genes as presented in Rashkovan et al (Rashkovan Cancer Discov 2022). Analyses were carried out on a subset of 4,871 metabolic genes obtained from the Reactome Pathway database (ID R-HSA-1430728). PROGENy scores were computed on the entire transcriptome as previously reported^36, 37^.

### Primary T-ALL cells cultures and transduction

T-ALL cells used in this study comprised patient-derived samples and low passage patient-derived NOD/SCID gamma c-/-(NSG) xenografts (1 or 2 passages). Fresh blasts were pre-stimulated for 3 days by cultivating them in RPMI-1640 medium supplemented with 20% Gibco serum (life technologies cat 10270106), cytokines (100 ng/mL stem cell factor, 100 ng/mL Flt3-ligand, 10ng/mL thrombopoietin) and insulin (20 nM) on a monolayer of OP9-DL1 stromal cells. Blasts were then collected and stromal cells eliminated. Subsequently, the leukemic blasts were plated on retronectin-coated wells and incubated with lentiviral vectors as previously described ^29^.

### Production and titration of lentiviral vectors

Pseudotyped lentiviral vectors used in this study were generated as previously described^29^ by transfecting 293T cells with a plasmid encoding the Gag-Pol proteins, a plasmid encoding BaEV glycoproteins and a plasmid carrying shRNA contracts and the GFP reporter. shRNA PTEN and control plasmids were a kind gift from Dominique Payet-Bornet and the shRNA ACLY plasmid and relative control shRNA were from Dharmacon. To determine the vectors titers, 293T cells were incubated overnight with serial dilutions of the vector supernatants. Upon replacing the medium with fresh DMEM, the cells were incubated for additional 3 days and the GFP^+^ cells was next determined by FACS analysis. The viral titers were also confirmed by Q-PCR, as previously described^29^.

### Western Blot

10^7^thymocytes were lysed for 30 minutes on ice with 100ul homemade HNTG Buffer (HEPES 50mM, NaCl 150mM, Glycerol 10%, Triton X-100 1%, 1,5mM MgCl_2_, 1mM EGTA) completed with a HALT protease & phosphatase inhibitor cocktail (thermofisher, Cat 78440). Protein samples were allowed to migrated on 4-20% Mini-PROTEAN® TGX™ Precast Gels (Bio-Rad Cat. 4561094) and then transferred on nitrocellulose membranes (Bio-Rad Cat. 1704158), which were subsequently incubated overnight at 4°C with the following primary antibodies: anti ATP citrate lysase antibody D1X6P (Cell Signaling 13390S), Anti Phospho ATP citrate Lysase antibody Ser455 (Cell Signaling 4331S), anti PTEN antibody (Cell Signaling 9188S), Anti Phospho-Akt Ser473 (D9E) XP® (Cell Signaling 4060S), Anti AKT antibody (Cell signaling 9272S) and anti-actin antibody (Abcam ab-14128). The revelation and analysis of the membranes were performed by ChemiDoc XRS+ (Biorad 1708265).

### Flow cytometric analysis

To stain for thymic subpopulations, the following murine antibodies were used: CD4 APC/Cy7 (Biolegend 100526), CD8 PE (ebiosciences 12-0081), TCRab Alexa647 (Thermofisher Scientific HM3621). To stain for intracellular antigens, murine thymocytes were first stained using CD4 and CD8 antibodies and then fixed and permeabilized using Fixation Buffer (Biolegend cat 420801) and Phosflow Perm Buffer III (cat 558050) solutions. Subsequently, the cells were stained overnight with primaries antibodies anti-pACLY (Abcam ab205430). Flow cytometric analyses were performed using a FACSAria™ III (BD Biosciences). Data were analyzed using FlowJo software.

### Targeted LC-MS metabolomics analyses

For metabolomic analysis the extraction solution was composed of 50% methanol, 30% acetonitrile (ACN) and 20% water. The volume of the extraction solution was adjusted to cell number (1 ml per 1E7 cells). After addition of extraction solution, samples were vortexed for 5 min at 4 °C and centrifuged at 16,000*g* for 15 min at 4 °C. The supernatants were collected and stored at −80 °C until analysis. LC/MS analyses were conducted on a QExactive Plus Orbitrap mass spectrometer equipped with an Ion Max source and a HESI II probe coupled to a Dionex UltiMate 3000 uHPLC system (Thermo). External mass calibration was performed using a standard calibration mixture every seven days, as recommended by the manufacturer. The 5 µl samples were injected onto a ZIC-pHILIC column (150 mm × 2.1 mm; i.d. 5 µm) with a guard column (20 mm × 2.1 mm; i.d. 5 µm) (Millipore) for LC separation. Buffer A was 20 mM ammonium carbonate, 0.1% ammonium hydroxide (pH 9.2), and buffer B was ACN. The chromatographic gradient was run at a flow rate of 0.200 µl min−1 as follows: 0–20 min, linear gradient from 80% to 20% of buffer B; 20–20.5 min, linear gradient from 20% to 80% of buffer B; 20.5–28 min, 80% buffer B. The mass spectrometer was operated in full scan, polarity switching mode with the spray voltage set to 2.5 kV and the heated capillary held at 320 °C. The sheath gas flow was set to 20 units, the auxiliary gas flow to 5 units and the sweep gas flow to 0 units. The metabolites were detected across a mass range of 75–1,000 *m*/*z* at a resolution of 35,000 (at 200 *m*/*z*) with the automatic gain control target at 106 and the maximum injection time at 250 ms. Lock masses were used to ensure mass accuracy below 5 ppm. Data were acquired with Thermo Xcalibur software (Thermo). The peak areas of metabolites were determined using Thermo TraceFinder software (Thermo), identified by the exact mass of each singly charged ion and by the known retention time on the HPLC column. Differential metabolite detection analyses were performed using the MetaboAnalyst software (version 5.0). Samples were normalized by median and data were log10-transformed and auto-scaled (Z-score) prior to data analysis. Normalization efficacy was verified by monitoring feature density distribution. Univariate analyses were carried out (either t-test or ANOVA based on the number of categories to compare) to identify statistically differentially detected metabolites (FDR < 0.05, absolute fold change >= 1.5). Partial least squares discriminant analyses were conducted to achieve multivariate dimensionality-reduction of the dataset. Hierarchical clustering analyses were realized using Euclidean distances and the complete clustering method using the top 50 deregulated metabolites obtained from t-test/ANOVA. Dendrograms indicate the relationship between samples, and metabolites separately. Enrichment analyses using MetaboAnalyst were carried out on differentially detected metabolites (FDR < 0.1, absolute fold change > 0.5) mapped to the KEGG database.

### Cholesterol quantification

Intracellular cholesterol levels were quantified using the Amplex Red Cholesterol Assay Kit (Invitrogen). Briefly, 2.10^6^ viable PDX blasts were collected from the indicated samples and washed with ice-cold PBS prior to testing according to the manufacturer’s instructions.

### Statistical analysis

Whereas otherwise indicated, significance levels of data were determined by Student’s t test, ANOVA or non-parametric equivalent tests depending on the normality of the data for the differences in mean values. Asterisks refer to the following p-values: * = p<0.05, ** = p<0.0, *** = p<0.001, **** = p<0.0001

## Acknowledgments

This work has been supported by a grant from Association Recherche sur le cancer (ARC) to MT. The authors wish to thank Ludovic Lhermitte for critically reading the manuscript and the animal facility technicians for animal husbandry.

## Supplementary figures

**Supplementary figure 1. ACLY mutant mice efficiently delete ACLY**

Representative western blot showing the levels of ACLY and PTEN in the thymus of PTEN^Δ/Δ^mutant, ACLY^Δ/Δ^mutant, PTEN^Δ/Δ^;ACLY^Δ/Δ^double mutant mice or PTEN^flox2^;ACLY^flox2^control animals.

**Supplementary figure 2. Reduced time to leukemia in PTEN^Δ/Δ^;ACLY ^Δ/Δ^double mutant mice**.

Time to leukemia in PTEN^Δ/Δ^ mutant mice and PTEN^Δ/Δ^;ACLY^Δ/Δ^ double mutant mice.

**Supplementary figure 3**. Double mutant mice do not develop T-ALL/lymphoma during 1 year follow-up.

Numbers of cells in the thymus **(A)**, spleen **(B)** lymph node cells **(C)** and bone marrow **(D)** of 1 1-year old PTEN^Δ/Δ^;ACLY^Δ/Δ^double mutant mice and PTEN^flox2^;ACLY^flox2^ control animals. Thymic populations **(E)** and percentages of CD4+ and CD8+ cells in the spleen **(F)**, lymph nodes **(G)** and bone marrow **(H)** and peripheral blood **(I)** of 1-year old PTEN^Δ/Δ^;ACLY^Δ/Δ^double mutant mice and PTEN^flox2^;ACLY^flox2^control animals. In mutant mice, carrying the YFP^flox2^reporter, thymic DN, DP, SP4 and SP8 and peripheral CD4+ and CD8+ cells are gated on YFP+ cells. Whereas otherwise indicated data are shown as average ± SE in at least 3 independent experiments, n=6 per group.

**Supplementary figure 4. Impact of ACLY deletion on lymphoid organs**.

Numbers of cells in the thymus **(A)**, spleen **(B)** lymph node cells **(C)** and bone marrow of 1-year old ACLY^Δ/Δ^double mutant mice and ACLY^flox2^control animals. Thymic populations **(D)** and percentages of CD4+ and CD8+ cells in the spleen, lymph node and bonne marrow and peripheral blood of 1-year old ACLY^Δ/Δ^double mutant mice and ACLY^flox2^control animals double mutant mice and PTEN^flox2^;ACLY^flox2^ control animals as evaluated by FACS analysis. In mutant mice, carrying the YFP^flox2^ reporter, thymic DN, DP, SP4 and SP8 and peripheral CD4+ and CD8+ cells are gated on YFP+ cells. Whereas otherwise indicated data are shown as average ± SE in at least 3 independent experiments, n=6 per group.

**Supplementary figure 5. Metabolomic analysis of PDX-derived T-ALL cells** Heatmaps of the most 50 deregulated metabolites in 8 PDX which underwent metabolic analysis **(A)**. Metabolic pathways enriched in PTEN altered patients **(B)** or PTEN GL patients **(C).**

