## Supplementary material for "ATP citrate lyase is an essential player of the metabolic rewiring induced by PTEN loss during T-ALL development": supplemenatry figures: E Supplementary figures.pdf

### Supplementary figure 1

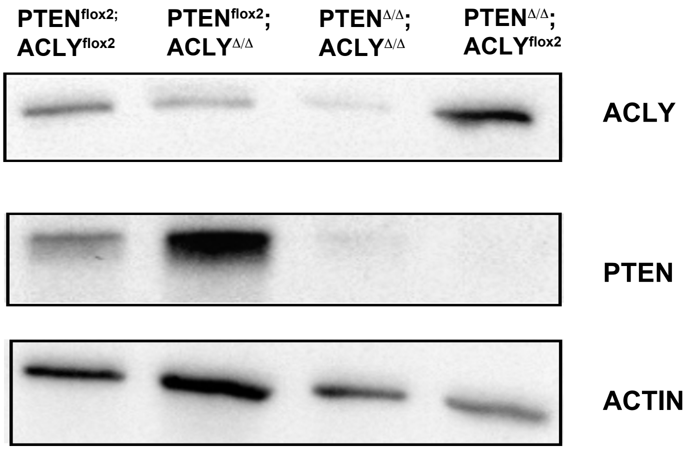

### Supplementary figure 2

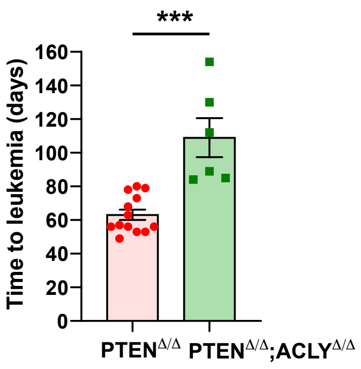

### Supplementary figure 3

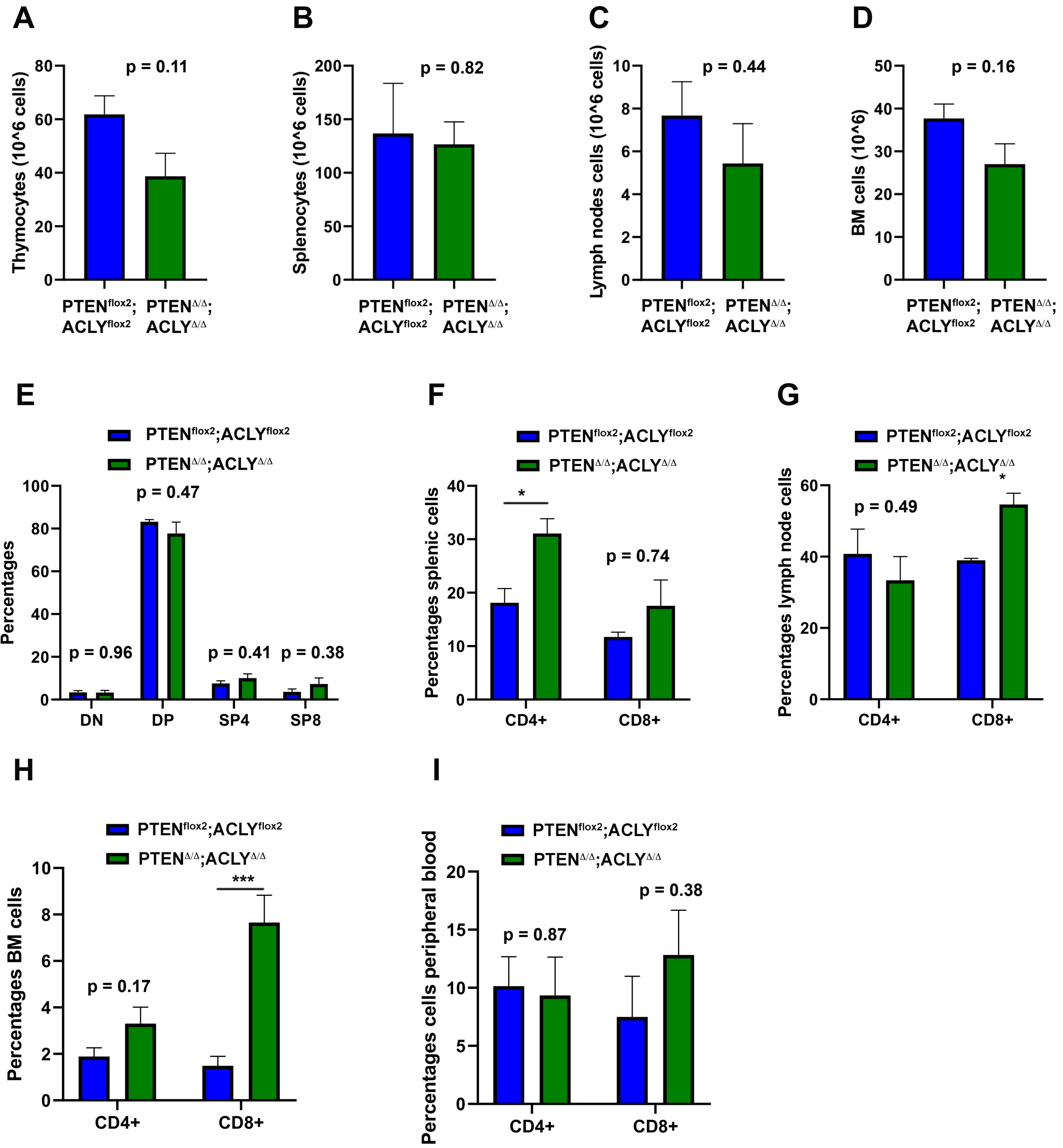

### Supplementary figure 4

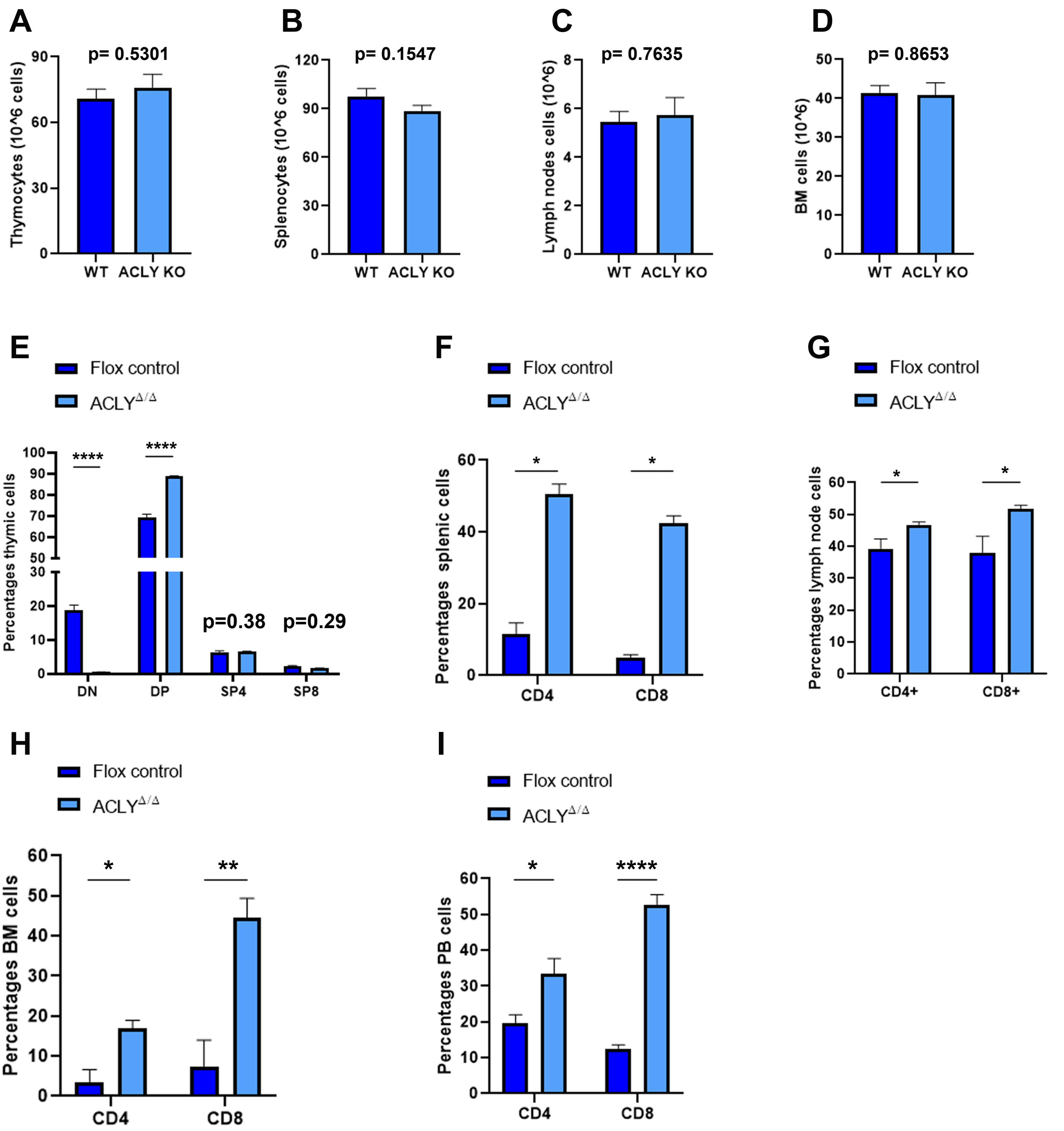

### Supplementary figure 5

A

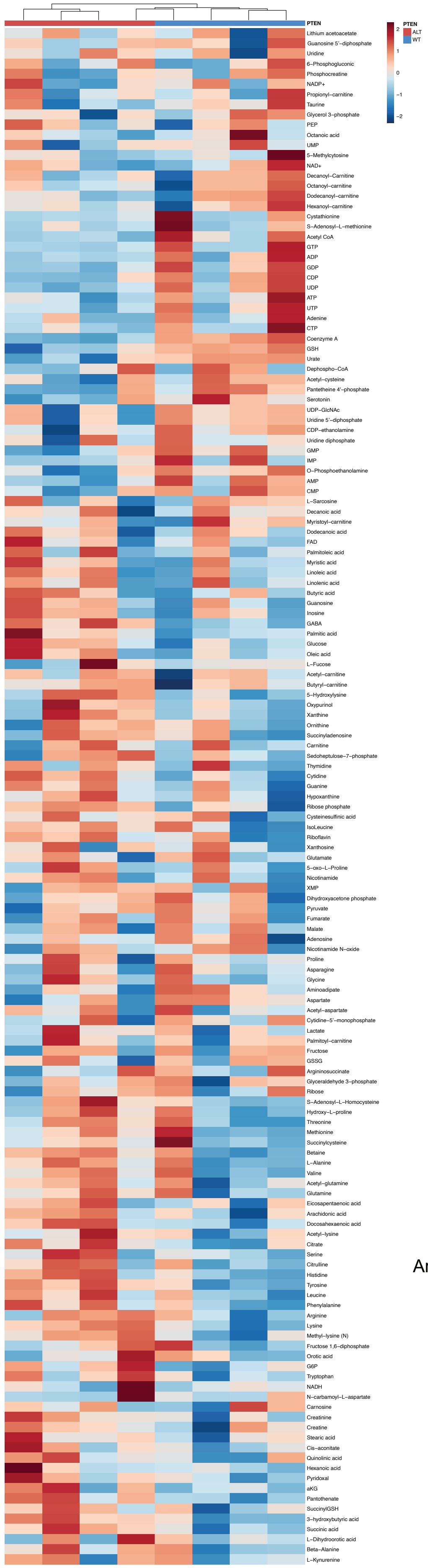


B

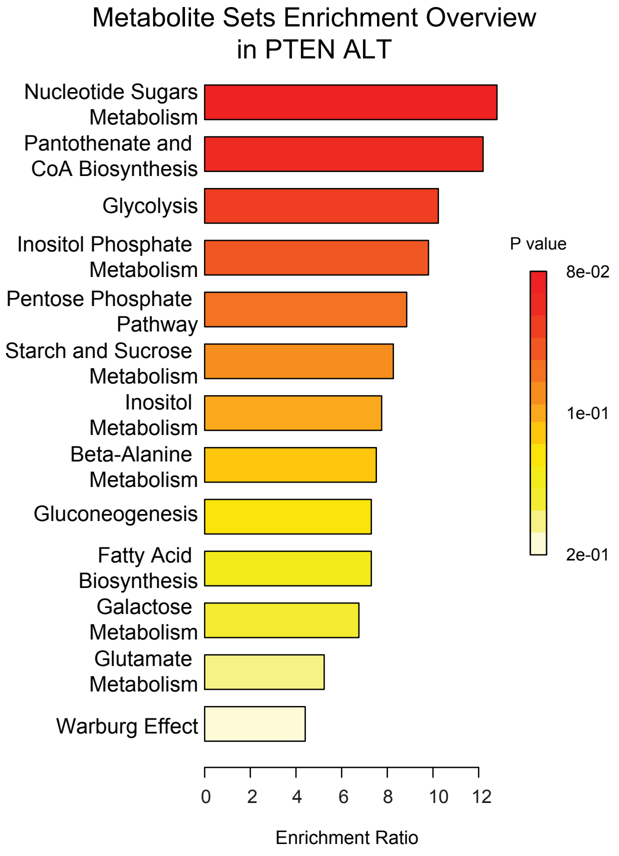


C

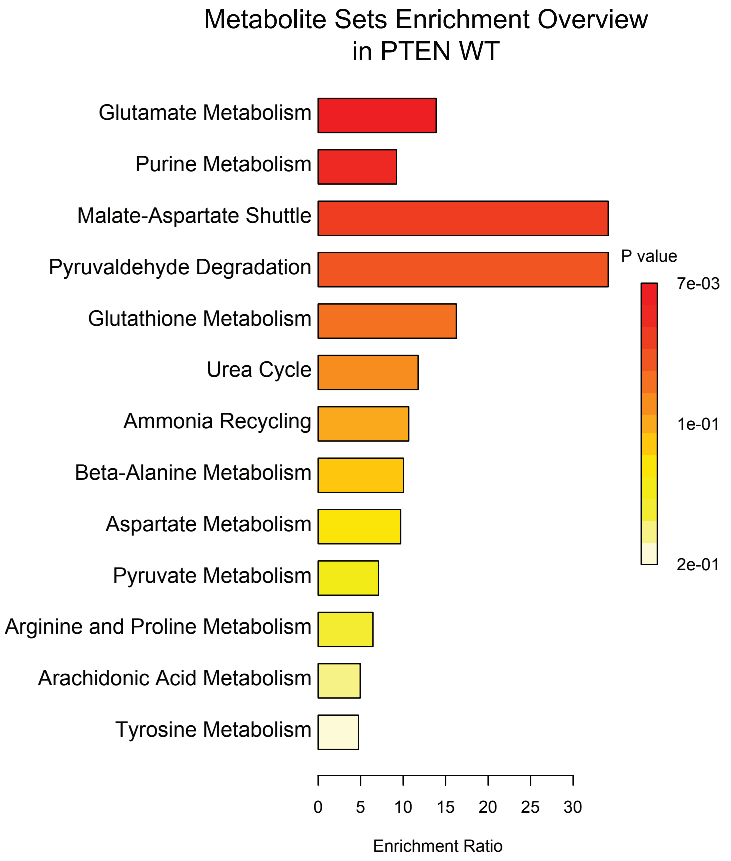
